# The *Staphylococcus aureus* Two-Component System AgrAC Displays Four Distinct Genomic Arrangements That Delineate Genomic Virulence Factor Signatures

**DOI:** 10.1101/277277

**Authors:** Kumari Sonal Choudhary, Nathan Mih, Jonathan Monk, Erol Kavvas, James T. Yurkovich, George Sakoulas, Bernhard O. Palsson

## Abstract

Two-component systems (TCSs) consist of a histidine kinase and a response regulator. Here, we evaluated the conservation of the AgrAC TCS among 149 completely sequenced *S. aureus* strains. It is composed of four genes: *agrBDCA*. We found that: *i)* AgrAC system (*agr*) was found in all but one of the 149 strains; *ii)* The *agr* positive strains were further classified into four *agr* types based on AgrD protein sequences, *iii)* the four *agr* types not only specified the chromosomal arrangement of the *agr* genes but also the sequence divergence of AgrC histidine kinase protein, which confers signal specificity, *iv)* the sequence divergence was reflected in distinct structural properties especially in the transmembrane region and second extracellular binding domain, and *v)* there was a strong correlation between the *agr* type and the virulence genomic profile of the organism. Taken together, these results demonstrate that bioinformatic analysis of the *agr* locus leads to a classification system that correlates with the presence of virulence factors and protein structural properties.

## Introduction

*Staphylococcus aureus* is a Gram positive human pathogen that has evolved considerable antimicrobial resistance during the clinical antibiotic era. It can cause a wide spectrum of infection types and severity, including soft tissue infection, bloodstream infections, pneumonia, osteomyelitis, and nosocomial device related infections (Yarwood & Schlievert, 2003; Lowy, 1998; Tong *et al*, 2015). The wide range of pathogenicity of *S. aureus* can be attributed to its ability to produce various secreted virulence factors, such as enterotoxin, hemolysin (*hla*), serine proteases (SspA), and TSST-1 (toxic shock syndrome toxin), as well as those mediating cell adhesion and host evasion (Nizet, 2007). Most of the virulence factors are regulated by a two-component system (TCS), AgrAC. AgrAC TCS is encoded by the *agr* locus and acts as a quorum sensing (QS) system in *S. aureus*. This QS mechanism modulates gene expression based on population density in response to environmental stimuli.

The *agr* locus of *S. aureus* contains four genes: *agrB, agrD, agrC* and *agrA*. *agrD* encodes for pre-peptide signal AgrD, which is modified to mature quorum sensing autoinducing peptide (AIP) and secreted to the extracellular space by the protein AgrB. *agrC* and *agrA* encode for histidine kinase and its cognate response regulator, respectively, and form the TCS. The *agr* operon can classify *S. aureus* strains into four variants or *agr* types (Dufour *et al*, 2002; Novick *et al*, 1995; Jarraud *et al*, 2000). These four variants (*agr* types) produce different AIPs that are characterized by different motifs of varying length. The AIP of one group has been identified to cross-inhibit the *agr* expression in the other groups (Jarraud *et al*, 2000; Shopsin *et al*, 2003; Geisinger *et al*, 2009). While much is known about the divergence of the *agr* variants (Dufour *et al*, 2002), most studies have focused on host proclivity (e.g., humans, bovine, etc.) (Lim *et al*, 2012; Jarraud *et al*, 2002; Wan *et al*, 2013; Schmidt *et al*, 2017) or on geographical location (Zaraket *et al*, 2007; Xie *et al*, 2011; Liu *et al*, 2012).

In this study, we predicted the *agr* type of all the completely sequenced *S. aureus* strains, irrespective of the host type or geographical location, and analyzed the divergence amongst the four *agr* types. We also proposed the evolutionary implications of these divergence.

## Results

The aim of this study was to compare the divergence of the four *agr* types in the strains of *S. aureus*. Therefore, we first predicted the *agr* types of the available completely sequenced strains and then subsequently set out to determine the *agr* type specific differences in i) the arrangement of genes in the *agr* locus, ii) sequence variation in the histidine kinase AgrC, and iii) the presence of different virulence genes.

### Agr types and ST typing

We analyzed 149 completely sequenced *S. aureus* strains from PATRIC database (Wattam *et* al, 2014) for the presence of *agr* locus. Of these, 148 strains were predicted to be positive for *agr* locus, based on the domain search of histidine kinase and response regulator proteins and querying the neighborhood genes for the presence of *agrB* and *agrD* (Methods and Materials). Out of these 148 Agr positive strains, 85, 42, 16, and 4 strains were predicted as strains belonging to Types I, II, III, and IV, respectively. This prediction was made by scanning the AgrD protein sequences by a custom python script (Supplemental Dataset: “determine_pheromoneType.py”) for the presence of conserved motif that are present in each AIP Type (Hawver et al., 2016). There was one exception to the type classification we observed. *S. aureus* strain TCH 959 did not have any conserved cysteine residue for AgrD and therefore could not be classified into any of the four types, thus, we classified it as “Type_undefined”.

In addition to the *agr* typing, Multilocus Sequence Typing (MLST) is used as a tool to further delineate strains according to the sequence similarity (Robinson *et al*, 2005). Although a specific *agr* type has been linked to specific MLST groups (Wright *et al*, 2005), MLST groups within a specific *agr* Type may show clonal divergence (Wright *et al*, 2005). Hence, we further classified strains within a *agr* Type based on MLST grouping.

MLST grouping of these 148 *S. aureus* strains were analyzed from the PATRIC database (Wattam *et al*, 2014) (Supplementary file S1). This analysis was unable to identify MLST for all the strains in our study. Hence, a manual literature curation was conducted to assign MLST groups to these unclassified *S. aureus* strains. The names of the strains and their types (*agr* and MLST) are provided in the supplementary file S1. The MLST grouping which we curated through manual literature search is highlighted in red in supplementary file S1.

### Orientation of *agr* genes classifies *S. aureus* strains into four specificity groups

Comparing the differences in the core gene arrangements within the *agr* locus would allow us to identify if the genomic features of the *agr* locus could contribute to the *agr* type specificity. To identify these differences, we analyzed the orientation of genes in the *agr* locus for each *agr* type. We found that the intergenic distances between genes in the *agr* locus differed among the four *agr* types (Figure 1) and were highly conserved within a specific *agr* type. For example, over 90% (77/85) of type I strains have a precise overlap of 3 bp between *agrB* and *agrD*, whereas genes are separated by 3 bp, 4 bp, and 2 bp in Types II, III, and IV, respectively. Similarly, the intergenic distance between *agrD* and *agrC* in type II was 30 bp, while the same distance was 25 bp in the other three *agr* types. The intergenic distance between *agrC* and *agrA* was conserved and separated by 19 bp in all the four types.

**Figure 1:**
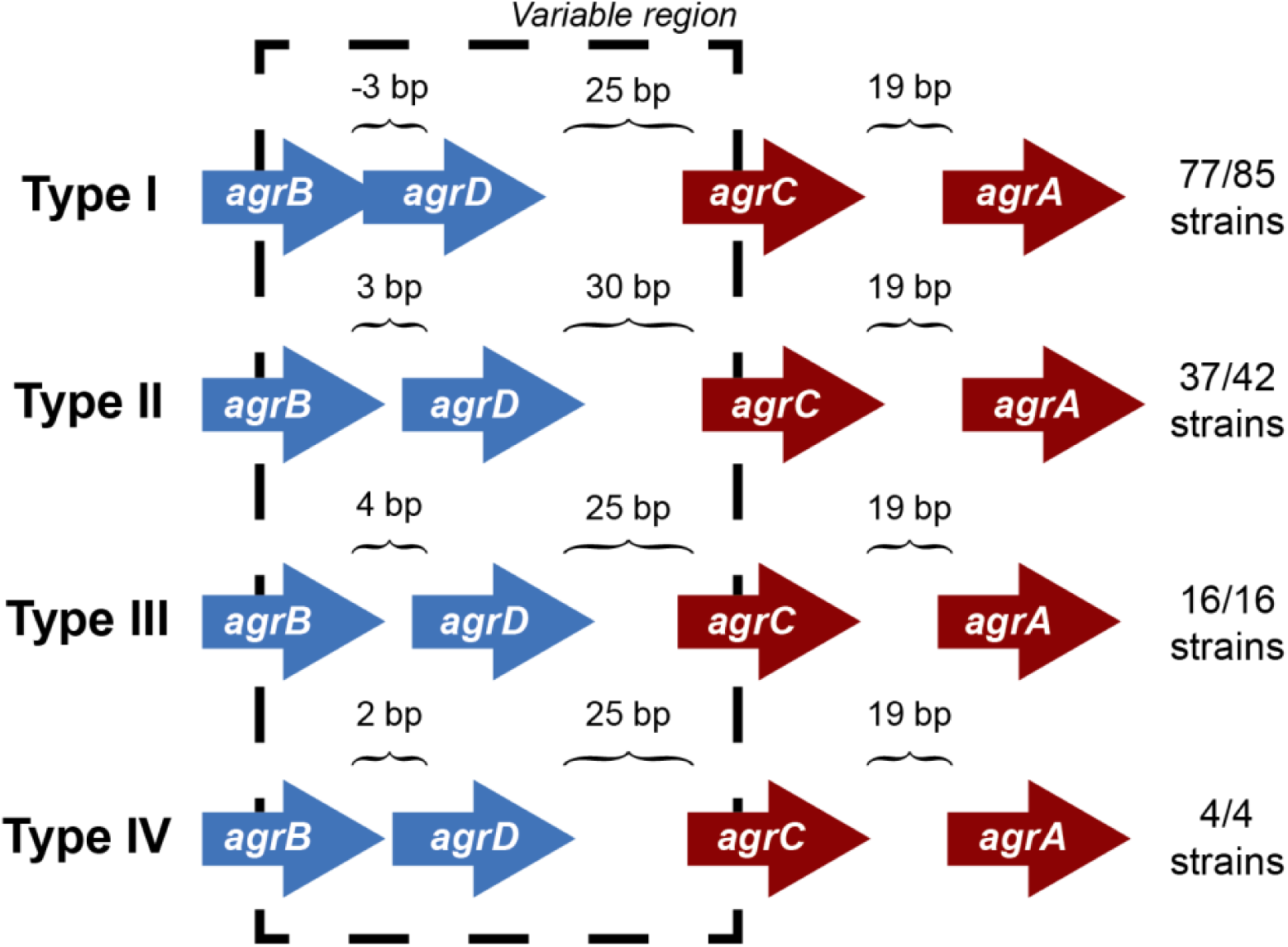
*agr* locus in different *agr* types. The intergenic distances between *agrB, agrD, agrC*, and *agrD* genes are conserved with *agr* types. The arrangement is true for 77 out of 85 strains in Type I, 37 out of 42 for Type II, and all the strains of type III and IV. The direction of arrows in the *agr* locus is the representation of those present in the positive strand. A similar arrangement occurs in the negative strand. The length of the arrows representing genes is not to scale. The black dotted line highlights the variable region in the *agr* locus.

However, we did observe a few non-canonical arrangements in types I (8/85) and II (5/42) which did not follow these same trends of intergenic distances indicated. Two of the type I strains (*S. aureus* strain T0131 and *S. aureus* strain HC1335) were found to contain a transposase gene between the hypothetical gene adjacent to *agrD* and *agrC* (Supplementary Figure S1). A previous study from Botelho *et al.*, identified IS256 transposase in these two strains and suggested that this transposase truncated *agrC* into two, which inhibited the gene and subsequently the regulation of *agr* locus [Botelho et al, 2016]. While we were unable to identify any conserved motifs in the hypothetical gene between *agrD* and *agrC* to define it as a histidine kinase, we did observe five transmembrane helices. This observation may suggest that the transposase cleaved *agrC* in a manner that separated the transmembrane domain from its catalytic domain, disrupting the regulation of *agr* locus. Nevertheless, even with the insertion of transposase or a hypothetical gene, type I maintained its canonical intergenic distances of -3, 25, and 19 bp between *agrB, agrD, agrC*, and *agrA*, respectively. The type II variants seem to deviate from the canonical intergenic distances of 3, 30, and 19 bp distance between *agrB, agrD, agrC*, and *agrA*, respectively, due to the insertion of hypothetical genes in the *agr* locus (Supplementary Figure S1). These variations identify sets that may be pseudo non-functional *agr* locus, as opposed to the canonical fully functional *agr* locus.

Thus, it was clear that the gene arrangements in the *agr* locus are specific to the *agr* types which may contribute to the type specificity of the AIP recognition and allow selective advantage over the niche selection.

### AgrC sequence variations among the *agr* types

In the previous section, we identified that the gene arrangements in the *agr* locus is specific to the *agr* types, suggesting that the AgrC protein sequences may show divergence according to the type of AIP it senses (Goerke *et al*, 2005; Ji *et al*, 1997; Robinson *et al*, 2005; Dufour *et al*, 2002; Geisinger *et al*, 2008). Thus, we further wanted to identify specific differences in all the AgrC protein sequences among the four *agr* types. The variance in AgrC proteins amongst *agr* types was identified in two ways: A) by investigating the overall differences in the biochemical content of the AgrC sequences among the four *agr* types and B) by scoping out individual amino acid variations between the types.

### A) Overall differences in the biochemical properties of AgrC proteins

We first looked at characterizing differences in the basic biochemical properties of AgrC among the four *agr* types, to give us a general idea of the changes occurring at the level of the entire protein sequence. For this, we wanted to see if AgrC protein sequences clustered together as described by these biochemical properties in a principal component analysis (PCA). The first two components (PC1-55% and PC2-34%) explained much of the variance seen. The first principal component (PC1) distinguishes between types I/IV, II, and III (Figure 2A). Type II proteins are enriched with a higher percentage of alanines, glycines, and leucines, with an overall greater molar percentage of small and basic residues and a higher isoelectric point (Figure 2B). In correspondence with these properties, type II proteins will also have a lower percentage of tryptophans, serines, glutamic acids, and isoleucines, with an overall lower molar percentage of acidic residues.

**Figure 2:**
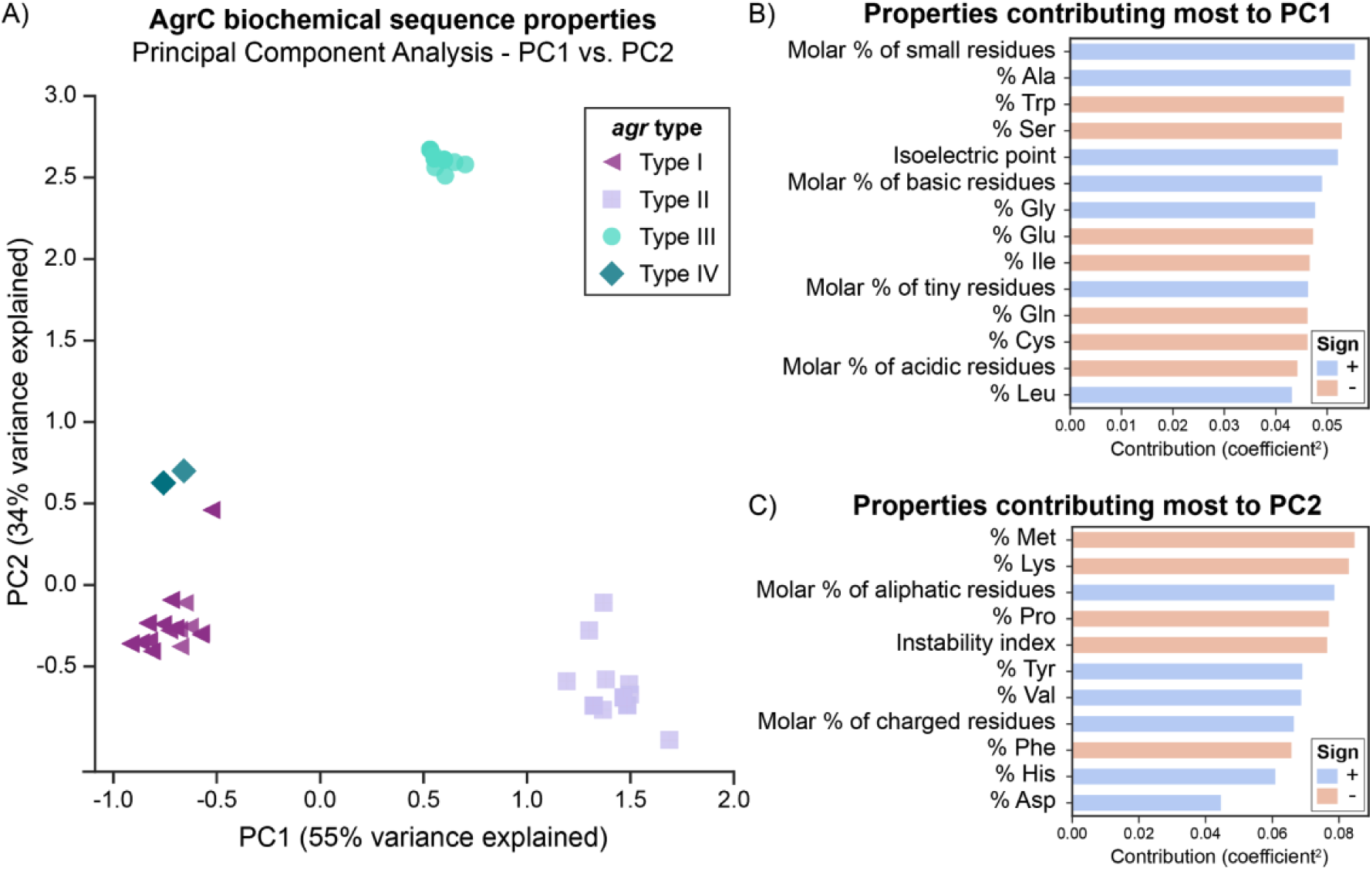
PCA of AgrC protein sequences across the *agr* types. A) A total of 89% of variance is explained with the first two components. Visually, the types cluster well together. A group of six type I strains (purple triangles) cluster close to type IV strains (dark green diamonds) and are commented upon in the main text. B) Properties that contribute most to principal component 1. A positive sign (blue bar) indicates that this property increases with PC1, while a negative sign (red bar) indicates that this property decreases with PC1. Properties shown are those with a contribution score greater than 1/(total number of properties). C) Properties that contribute most to principal component 2.

Six Type I AgrC proteins (purple triangles in Figure 2A) belonging to ST59 Type strains clustered near the Type IV AgrC proteins (dark green diamonds), potentially suggesting that Type IV may have diverged from these Type I strains during evolution. Goerke et al have also noted a subgroup of type I which was related to Type IV (Goerke *et al*, 2005). The third principal component (Supplemental Figure S2) explains an additional 5% of variance, and interestingly clusters these six strains alongside the type IV strains. Thus, from this component, these strains are characterized by an enrichment of polar residues, particularly histidine, with a lower molar percentage of non-polar residues. Finally, the second principal component (PC2) (Figure 2C) distinguished type III (cyan circles) very well from the other types and is characterized by higher molar percentage of aliphatic amino acids, charged amino acids, and valines, along with a lower percentage of methionines, lysines, prolines, phenylalanines, and a lower instability index (predicted to be more stable). We also attempted PCA on each of the domains of AgrC (extracellular or intracellular loops, transmembrane helices, or cytoplasmic domain), with the results presented in Supplemental Figure S3. Interestingly, we saw the same clustering of the types for most of the sub-domains.

### B) Amino acid variants in AgrC proteins

From the previous section, it was clear that the biochemical properties of the AgrC protein was specific to the *agr* type and a few strains of type I may form a subgroup closer to type IV. We further looked at the specific amino acid variations in the AgrC protein that may have contributed to the distinctive variations in the biochemical properties among the four *agr* types. For this analysis, we compared the protein sequences of AgrC for each *agr* Type against a type I AgrC (AgrC-I) from *S. aureus* (strain COL) as a reference sequence (UniProt: A0A0H2WWL2) (Srivastava *et al*, 2014). This comparison was limited to AgrC proteins forming canonical gene arrangements, as those in non-canonical arrangements were mostly truncated by insertion sequences and hence amino acid variations would not be correctly identified. Furthermore, the amino acid variations observed in each *agr* type were mapped onto the predicted six transmembrane domain model (Lina *et al*, 1998) of the reference AgrC-I sequence A0A0H2WWL2 and were classified using Grantham scores to rate their changes on a scale of “conservative” to “radical” (see Materials and Methods).

Only a few strains of type I deviated from the reference AgrC-I sequence and, moreover, these variations were mostly conservative in nature, demonstrating that the change in the amino acids within the Type I would hypothetically not affect the AgrC structure (Figure 3 and Figure S4). The variations observed within Type I strains were seen to be dictated by the MLST typing of the strains (Supplementary file S2), e.g., the mutation L3T occurred in six strains of type I belonging to the ST59 group. A few notable changes included a mutation of V42G which appeared in 22 strains of Type I (ST7, ST72, ST398, ST25, ST464) in the first extracellular loop, and F114S which appeared in only two related strains (ST25) near the predicted second extracellular loop (Figure 3D, Supplemental Figure S4). The AgrC sequences for types II and III strains were found to be the most divergent from the reference AgrC-I (Figure 3A). The amino acid variations were mainly concentrated on the first 200 residues, which comprise the N-terminal transmembrane region responsible for sensing AIPs (Figure 3B). We determined that in types II and III, the majority of amino acid variations were from nonpolar to nonpolar residues (Figure 3C), likely to preserve the helical transmembrane structure as they were mostly found in these regions. Some of the amino acid variations observed in type II were classified as radical changes (Figure 3E, Supplemental Figure S5). Specifically, a few biochemically divergent residues (e.g., G29F, S64F, F65S, C91L, marked in red in Figure 3E) were observed, all of which may rearrange the three-dimensional orientation of the transmembrane helices and consequently their helix-helix contacts in type II strains. In type III strains, none of variations were classified as radical changes (Supplemental Figure S6). type IV showed very little divergence compared to type I (Supplemental Figure S7). The amino acid variations in the transmembrane as well as catalytic domains for each Type are shown in Supplemental Figures S4-S7.

**Figure 3:**
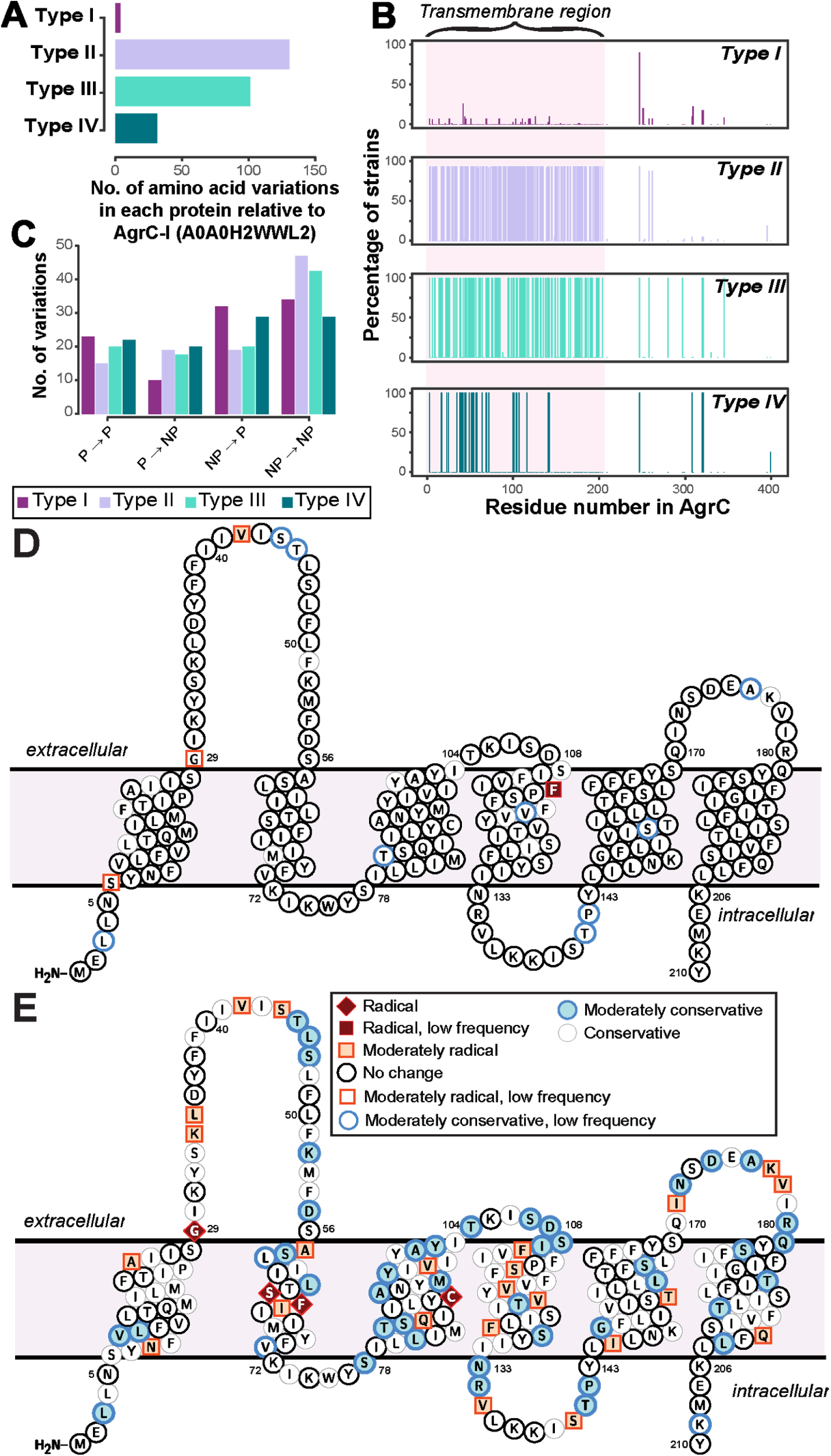
Sequence divergence in AgrC of four *agr* types. **A)** The number of amino acid variations in each protein of *agr* types. type I shows the least variation, as the reference strain used for the analysis is of type I. AgrC sequence of type II and type III have the highest number of amino acid variations. **B)** The percent of strains having a variation in that residue number. Around 92% and 100% of strains of type II and type III, respectively, have variations mostly in the first 200 residues of AgrC protein. **C)** Properties of amino acid variations in each type show that the majority of variations are between non-polar amino acids in type II and type III, meaning there is a much higher divergence in the transmembrane region. Abbreviations: NP, non-polar; P, Polar. D, E: specific amino acid divergence between type I and type II. The predicted transmembrane topology of AgrC-I reference sequence from CCTOP with highlighted amino acid residues that tend to diverge in **D)** type I strains and **E)** type II strains, from this study. The color coding is per biochemical and biophysical property of the amino acid residue mutated. Radical, moderate, and conservative is the nomenclature given in the order of amino acid divergence. Radical is a vast difference between amino acid properties and could change the properties of the protein. Conservative is an amino acid substitution which can be tolerated by the protein. Moderate is a biochemically moderate change in the amino acid substitution. Low frequency: amino acid variations that occurred only in very few strains.

Finally, we inspected variations which occurred together (referred to as co-occurring mutations) in the cytoplasmic domain, of which the CA domain has been crystallized (Srivastava *et* al, 2014) to understand the stability of the AgrC proteins for each Type. The mutation of P247 to a threonine corresponds to a phosphotransfer specificity residue (A268) found in a previous study of a similar two-component system (Podgornaia *et al*, 2013) and does not seem to be a type specific mutation itself. This mutation was found to commonly co-occur with additional mutations at S320 and S321 which are located distal to the ATP-binding domain (Supplemental Figure S8). However, we were able to observe various combinations of co-occurring variations that were type specific, e.g., a set of seven type I AgrC proteins had the co-occurring mutations of P247T, Y251F, S320T, S321R, and T345S. The downstream effects of these mutations remain difficult to elucidate even with experimental data (Capra *et al*, 2010; Podgornaia *et al*, 2013) but likely have impacts on both the specificity and strength of binding to cognate receptors. We inspected the impact of these co-occurring mutations on the stability of the AgrC protein by using a homology model of the dimeric cytoplasmic domain, and each set of the co-occurring mutation was predicted to be significantly destabilizing (Supplemental Table ST1). This destabilization suggests that these mutations may increase the propensity of the kinase to seek out its binding partners (ATP or a response regulator), then induce additional phosphotransfer reactions and lower response times to environmental stimuli. Another potential consequence would be an increase in the likelihood of non-cognate receptors to bind (Srivastava *et al*, 2014; Studer *et al*, 2013).

### Comparison of presence of virulence genes in the four specificity *agr* types

To further determine whether there is a correlation between the *agr* types and the presence/absence of the virulence factors, we evaluated the presence of virulence genes in all the 148 *agr* positive strains.

We predicted 216 virulence genes in total to be present in the 148 *agr* positive strains. Of those predicted, 104 were present in all 148 strains (Figure 5A). Some of the conserved virulence genes were capsular polysaccharide genes (*cap5A-G* genes), aureolysin, autolysin, gamma, and beta hemolysin (*hlg, hlb*), genes encoding iron regulated proteins (*isd* genes), and staphylococcal protein A (*spa*). *S. aureus* strain SA564 was predicted to have the highest number of virulence genes (189 out of 216) and a mecA negative strain of *S. aureus* strain SA17_S6 was found to have the least number of virulence genes (131 out of 216) (Supplementary file S3). Although most of the virulence genes seemed to be conserved in all the strains of *S. aureus* (Figure 5A), a classification tree of presence and absence of virulence genes suggested a few differences among the types (Figure 5B). For example, strains in which *set15* exotoxin is absent and *set30* exotoxin and *hld* are present may be characterized as Type I strains. Similarly, Type III can be characterized by the absence of *set15, set30*, and the presence of *cap8H*.

**Figure 5:**
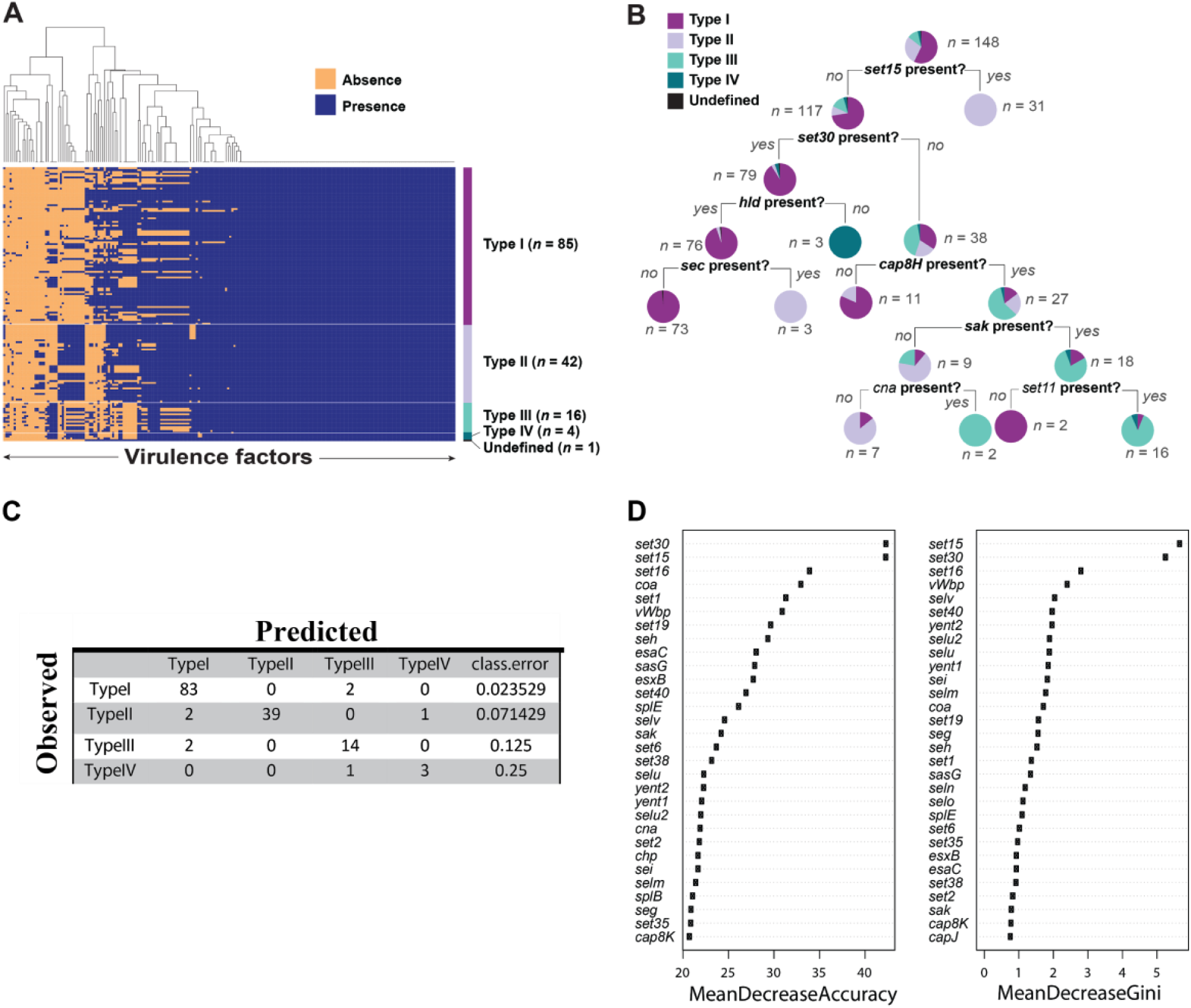
Virulence factors in *S. aureus* strains. A) Heat map showing presence and absence of virulence genes in each strain of *S. aureus*. Red depicts absence and green depicts the presence of a virulence genes. White line demarcates the heatmap into *agr* types. B) Classification tree reveals differential regulation of various virulence genes in *S. aureus* dictated by *agr* types. C) Accuracy of the random forest model in predicting *agr* types. D) The variable importance computed by random forest algorithm.

Next, we built a predictive model using random forest classifier to classify *agr* type based on the virulence phenotype. Random forest is a machine learning technique which builds an ensemble of classification trees and then takes the average of all the predictions to determine the class. The model showed an accuracy of 94.5% with an error rate of 5.44% in predicting the *agr* types. Of 85 Type I strains, 97.6% were correctly predicted. Similarly, 92.8% of type II, 87.5% of type III, and 75% of type IV were correctly predicted by this model (Figure 5C). The high accuracy of the model to correctly predict the *agr* type of the strains based on virulence genes demonstrates that virulence genes can be used to discriminate *agr* types. Similar to the decision tree shown in Figure 5B, we identified virulence genes that are important for accurate prediction of the *agr* types (Figure 5D). Random forest algorithm allows two Types of importance measurements: ‘mean decrease in accuracy’ and ‘mean decrease in gini coefficient’. The mean decrease in accuracy measures how the accuracy of the model decreases if a variable is dropped. The higher the decrease in accuracy due to exclusion of a variable, the more important that variable is considered. The mean decrease in gini coefficient measures the contribution of the variable towards the homogeneity of the nodes in the random forest tree. We utilized the variable importance measure in random forest algorithm to identify important virulence genes.

With these results it was evident that the presence or absence of virulence factors are dependent on the *agr* type of the *S. aureus* strains.

## Discussion

In this study, we analyzed the divergence among the *agr* types of *S. aureus* with respect to the genomic features and the virulence capacity and found that the genomic features of the *agr* locus and the virulence capacity of the *S. aureus* are strongly correlated to the *agr* types.

We identified that the gene arrangement of the *agr* locus is specific to the particular *agr* types. However, we also identified some strains of type I and type II which may have incorporated a “self-destruct” mechanism of the *agr* inactivation by incorporating transposase or additional hypothetical genes in the *agr* locus. Inactivation of *agr* appears to confer a fitness advantage under the conditions of antibiotic selection, *in vivo* environment, and biofilm formation (Botelho *et al*, 2016; Paulander *et* al, 2012; Tan *et al*, 2015). Loss of *agr* function has been associated with resistance to host cationic antimicrobial peptides, tolerance to vancomycin, and development of intermediate resistance to vancomycin, a phenotype that is associated with treatment failure in bloodstream infections (Fischer et al., 2011; Mwangi et al., 2007; Sakoulas et al., 2002, 2005; Tsuji et al., 2007; Vandamme et al., 1997). Therefore, the subsequent study of *agr* function from isolates derived from healthcare settings as compared to those from the community, as well as *agr* function with respect to site of infection, may shed further insights on the role of *agr* inactivation in the pathogenesis of different infections.

We also identified the specific biochemical properties and the amino acid variations that distinguished AgrC proteins among the four *agr* types. The biochemical properties of the overall protein as well as the sub-domains were found to be type specific, highlighting the fact that individual domains contribute to the structure-function relationship of AgrC. We also observed that the variant *agr* alleles may form within the same *agr* types depending on the MLST typing, however, this phenomenon was more evident in AgrC type I strains. Strains belonging to ST59 had different biochemical properties and amino acid variations than the canonical *agr* type I strains. We observed that ST59 strains were more related to type IV strains, however they had not attained a full transformation to type IV strains. A similar evolutionary model was proposed by Robinson et al (Robinson *et al*, 2005) where they hypothesized that amino acid variations may occur in the *agr* locus which may affect *agr* activity beyond the specificity of the four *agr* types. The amino acid variations did seem to be concentrated in the transmembrane domain of the AgrC protein, but we could also identify specific co-occurring mutations in the catalytic domain that were type specific. We predicted that these mutations can lead to protein destabilization and change the function of the protein either by increasing the catalytic activity or by decreasing the substrate specificity (Studer *et al*, 2013; Srivastava *et al*, 2014).

Finally, we predicted *agr* type specific production of virulence genes in *S. aureus* strains (Figure 5). This distinction may be because different *agr* type strains secrete and recognize different AIP signals and this inherent variability to sense diverse signals correlates with pathogenesis (Yarwood & Schlievert, 2003). In addition, random forest classifier enabled us to correctly predict the *agr* type with an accuracy of 94.5%. The accuracy of the model confirmed our hypothesis that the virulence capacity is indeed correlated with the *agr* type of the strain which may give them a more competitive advantage over others. It is important to note that although ST59 strains (type I strains) have diverged from the rest of the type I strains, as seen from the amino acid variations in the AgrC sequence, the divergence is not enough to differentiate the virulence capacity of these strains.

In conclusion, our study suggests that during evolution, *agr* typing and the gene arrangements of the *agr* locus may have evolved together, giving rise to differential virulence capacity. The AgrC sequences subsequently diverged, giving rise to variant *agr* alleles among the same *agr* types owing to environmental pressure and competitive advantage.

## Methods

### *S. aureus* genomic data

The genomic sequences of 149 completely sequenced *S. aureus* strains were downloaded from the PATRIC database (sequenced until December 2016) (Gillespie et al., 2011) and re-annotated them with the Prokka v1.12 (Seemann, 2014) annotation tool to identify protein coding genes. The re-annotation was performed to standardize annotations across all genomes. Draft genomes were excluded from the analysis because of their incompleteness.

### *In silico* prediction of the *agr* locus

The protein concatenated fasta files of all the acquired 149 strains of *S. aureus* obtained from Prokka were searched for the presence of the domains of histidine kinases and response regulators using HMMsearch from HMMER package (Eddy, 1998). Hidden Markov Model (HMM) profiles of the histidine kinases and response regulators from Pfam database were utilized to scan the protein sequences. An e-value of 0.01 and score >= 25 was taken as a threshold to filter the hits from HMMsearch. For identifying the *agr* operon, neighborhood genes were scanned for the presence of *agrB* and *agrD* genes from the protein feature table obtained from Prokka.

### Classification of strains based on AgrD and detection of chromosomal arrangement in *agr* Types

Based on the four Types of AIPs in *S. aureus*, we classified *S. aureus* strains into four *agr* types. Protein sequences of AgrD for all the *S. aureus* strains were scanned for the presence of the conserved motifs that are present in each AIP by a custom python script “determine_pheromoneType.py”. The corresponding strain was accordingly grouped into *agr* types. Further, the differences in the chromosomal arrangement of genes in the *agr* locus for each *agr* type were investigated. The chromosomal arrangement was identified in the form of “gene context”, defined by conserved intergenic distances between the genes in the *agr* locus.

### AgrC sequence variations among the *agr* Types

To analyze sequence variation in AgrC across Types, we utilized the ssbio package to inspect the biochemical properties of all sequences as well as mutations when aligned to a reference sequence (Mih *et al*, 2018). All strain sequences were loaded and those of non-standard length (∼430 residues) were ignored for this analysis. For the principal component analysis (PCA), we calculated general biochemical properties of the full sequences using the Biopython ProtParam and EMBOSS pepstats tools (Rice *et al*, 2000; Cock *et al*, 2009). These gave general descriptors such as the percentage of polar, non-polar, aromatic, small amino acids, etc. We used these descriptors to create a feature matrix which was then normalized with the Python scikit-learn package (Pedregosa *et al*, 2011). This feature matrix was then used for PCA, and the largest contributors to the first three principal components were then analyzed.

We conducted pairwise sequence alignments using the EMBOSS needle tool with default parameters (Rice *et al*, 2000) of all AgrC Types to the reference AgrC sequence, set to the available crystallized protein of the cytoplasmic domain (UniProt: A0A0H2WWL2, PDB: 4BXI) (Srivastava et al., 2014). The reference sequence is a type I AgrC protein of *S. aureus* strain COL. We then analyzed the sequence diversity among types with respect to the reference sequence in terms of amino acid variations at each residue. Variations were scored by the Grantham score, which is a simple measure of biochemical and biophysical property differences of amino acids (Grantham, 1974). These scores of variations for each AgrC type, along with the frequencies of variations observed in each strain types, were plotted on the 6 transmembrane model of AgrC-I (Lina *et al*, 1998), using the Protter web service (Omasits *et al*, 2014). Variation within the cytoplasmic domain was additionally mapped to a homology model from SWISS-MODEL (template PDB: 4JAU) as well as the crystal structure (PDB: 4BXI). Stability changes incurred from these mutations were predicted using FoldX and the included BuildModel function (Schymkowitz *et al*, 2005; Guerois *et al*, 2002).

### Virulence genes prediction and decision tree

We downloaded experimentally studied virulence genes in *S. aureus* from virulence factor database (VFDB) (Chen *et al*, 2005) and did a Blast search on the protein fasta sequences of all the studied *S. aureus* strains (Altschul *et al*, 1990) to predict potential virulence genes in *S. aureus* strains. A threshold of 85% identity and E-value 0.001 was set to only identity significant hits. A decision tree was constructed using rpart package in R based on presence/absence of virulence genes in strains (Therneau et al., 2017).

### Classification of *agr* types based on virulence genes

We built random forest predictive model using virulence factors as features to classify samples by *agr* type using the “randomForest” package in R (Liaw et al., 2015). Random forest classifier was built by growing 5000 trees. For detecting important virulence genes, we used variable importance measure in “randomForest” algorithm.

## Acknowledgements

We would like to thank Dr. Ke Chen for helpful discussion. We would also like to thank Marc Abrams for proofreading and editing of the manuscript.

